# Heterotrimeric G Protein–RasGAP Coupling Drives Adaptation During Chemotaxis

**DOI:** 10.64898/2026.02.24.707728

**Authors:** Xuehua Xu, Riley Kim, Haneul Hyun, Ranti Dev Shukla, Tian Jin

## Abstract

Chemotaxis enables eukaryotic cells to detect and migrate along extracellular chemoattractant gradients spanning several orders of magnitude. This remarkable dynamic range relies on adaptation, a process that allows cells to reset their signaling machinery while preserving sensitivity to incremental changes in stimulus intensity. Although numerous actin-dependent feedback mechanisms have been characterized, the molecular basis of adaptation within the actin-independent core gradient-sensing module remains poorly understood. Here we identify the Ras GTPase-activating protein C2GAP1 as a critical F-actin–independent effector of the heterotrimeric G protein Gα2 in *Dictyostelium discoideum*. Using cytoskeleton-free gradient-sensing cells, quantitative imaging, biochemical assays, FRET-based G-protein activation measurements, and structural modeling, we demonstrate that C2GAP1 controls concentration-dependent adaptation during gradient sensing. Mechanistically, C2GAP1 directly associates with Gα2 in both GDP- and GTP-bound states, with preferential binding to activated Gα2, thereby sustaining membrane recruitment and locally attenuating signaling. Loss of C2GAP1 enhances G-protein activation, disrupts front-specific inhibition, and impairs rapid reorientation in dynamic gradients. These findings define a direct coupling between heterotrimeric G proteins and a RasGAP as a core adaptive module that calibrates gradient sensing across wide concentration ranges.

**Highlights:** Chemotaxis, the directional migration of cells along chemoattractant gradients, underlies processes such as neuron patterning, lymphocyte recruitment, cancer metastasis, and *Dictyostelium discoideum* development. The hallmark of eukaryotic chemotaxis is the ability to sense and respond to gradients spanning wide concentration ranges through cellular adaptation. This process involves three interconnected modules: gradient sensing, cell polarity, and migration, with gradient sensing as the foundation. While many components of GPCR-mediated signaling are known, the molecular mechanisms driving adaptation remain unclear. Here, we show that the heterotrimeric G protein α subunit interacts with RasGAP C2GAP1 to mediate adaptation during gradient sensing and chemotaxis.

## INTRODUCTION

Chemotaxis is a directional cell migration guided by chemoattractant gradients. This cellular behavior plays essential roles in numerous physiological processes, such as neuron patterning, recruitment of lymphocytes, angiogenesis, metastasis of cancer cells (Condeelis et al., 2005; Murphy, 1994; Parent et al., 1998). The social amoeba *Dictyostelium discoideum* has served as powerful model system for dissecting the conserved signaling architecture of eukaryotic chemotaxis (Jin et al., 2008; Van Haastert and Devreotes, 2004). In this system, chemotactic responses to cyclic AMP (cAMP) are mediated by the G-protein–coupled receptor cAMP receptor 1 (cAR1), which couples to the heterotrimeric G protein composed of Gα2 and Gβγ subunits (Parent and Devreotes, 1999). Upon ligand binding, cAR1 activates heterotrimeric G proteins through dissociation of Gα2 from Gβγ, thereby initiating downstream signaling cascades that drive directed migration (Janetopoulos et al., 2001; Jin et al., 2000; Xu et al., 2010). The molecular mechanisms governing cAR1/Gα2Gβγ-mediated chemotaxis have been extensively investigated. Downstream of G-protein activation, Ras proteins serve as central signaling nodes that activate multiple pathways—PI3K, TORC2, PLA2, ElmoE, and soluble guanylyl cyclase (sGC)—which together coordinate chemotactic signaling and cell movement (Cai et al., 2010; Charest et al., 2010; Funamoto et al., 2002; Iijima and Devreotes, 2002; Yan et al., 2012).

Several regulatory mechanisms acting at the level of G proteins have been described. Ligand-induced phosphorylation of cAR1 reduces its coupling to heterotrimeric G proteins and functions as a receptor desensitization mechanism (Caterina et al., 1995; Hereld et al., 1994; Xiao et al., 1999). G-protein–interacting protein 1 (GIP1) binds Gα2 and facilitates its shuttling between the cytosol and plasma membrane, ensuring sufficient availability of G proteins at the membrane during chemotaxis (Kamimura et al., 2016). ElmoE is an effector of Gβ that associates with Gβ to activate Rac and thereby promote cell migration (Yan et al., 2012). In addition, the nonreceptor guanine nucleotide exchange factor Ric8 acts as a Gα2 effector to amplify Gα2 signaling (Kataria et al., 2013). Importantly, chemotaxis is a coordinated process comprising three conceptually distinct but interconnected modules: gradient sensing, cell polarity, and cell migration. Gradient sensing can be uncoupled from both initial polarity establishment and actin-based motility (Parent and Devreotes, 1999). For example, cells treated with actin polymerization inhibitors lose their filamentous actin–based cytoskeleton and become immobile; nevertheless, they retain the ability to sense a chemoattractant gradient. This observation demonstrates that the core gradient-sensing machinery operates independently of F-actin–based cytoskeletal structures. In contrast, the functions of ElmoE and Ric8 require an intact actin cytoskeleton, indicating that these regulators operate within F-actin–dependent feedback loops rather than within the actin-independent core gradient-sensing module. Thus, despite extensive characterization of actin-dependent signaling pathways, the key components that constitute the actin-independent core machinery responsible for gradient sensing remain largely unidentified and represent a critical gap in our understanding of eukaryotic chemotaxis.

Chemotactic cells detect and respond to an enormous concentration range of chemoattractants. For example, *D. discoideum* cells chemotax toward their chemoattractant cAMP gradients from 10^-9^ to 10^-5^ M (Janssens and Van Haastert, 1987). To chemotax through gradients with such a large concentration range, cells employ a mechanism called adaptation (Parent and Devreotes, 1999). The temporal properties of adaptation were first characterized using uniformly applied chemoattractant stimuli (Dinauer et al., 1980; Van Haastert, 1987a). In response to sustained stimuli, cells exhibit a transient signaling response that subsequently returns toward baseline despite the continued presence of the stimulus, a phenomenon of adaptation. A defining feature of adaptation is that cells become insensitive to the persistent stimulus while retaining the ability to respond to further increases in chemoattractant concentration. In *D. discoideum*, cAR1 GPCR (cAMP receptor)-mediated phosphatidylinositol (3,4,5)-trisphosphate (PIP_3_) responses display all hallmark features of chemoattractant sensing and adaptation (Parent and Devreotes, 1999). When cAMP is applied uniformly, the signaling pathway leading to PIP_3_ production can be divided into four sequential steps with distinct kinetics. First, cAMP binds to cAR1 receptor (Klein et al., 1988; Van Haastert, 1987b). Second, activated cAR1 induces a persistent dissociation/activation of heterotrimeric G-proteins (Gα2Gβγ) (Janetopoulos et al., 2001; Jin et al., 2000), indicating that adaptation occurs downstream of G protein activation. Third, Ras proteins are activated by guanine nucleotide exchange factors (GEFs), which catalyze the exchange of Ras-GDP for Ras-GTP; Ras-GTP is subsequently inactivated by Ras GTPase-activating proteins (RasGAPs), which stimulate its intrinsic GTPase activity (Charest et al., 2010; Insall et al., 1996; Kortholt et al., 2013; Sasaki et al., 2004). Uniform cAMP stimulation elicits a transient Ras activation followed by concentration-dependent, imperfect adaptation (Xu et al., 2022b; Xu et al., 2017). Fourth, Ras-activated PI_3_K phosphorylates phosphatidylinositol 4,5-bisphosphate (PtdIns(4,5)P2, PIP_2_) to phosphatidylinositol (3,4,5)-trisphosphate (PtdIns(3,4,5)P3, PIP_3_) in the membrane; while lipid phosphatase PTEN transiently dissociates from the membrane to permit accumulation of PIP_3_ and then returns to dephosphorylate PIP_3_ back to PIP_2_ (Funamoto et al., 2002; Iijima and Devreotes, 2002). Collectively, those observations identify Ras activation as the earliest step in the GPCR-mediated signaling cascade that exhibits adaptive behavior. The *D. discoideum* genome encodes at least 18 putative RasGAP proteins, suggesting extensive regulatory potential at the level of Ras inactivation. To date, two RasGAPs, NF1 (*nf1*) and C2GAP1 (*c2gapA*), have been shown to play critical roles in adaptation and chemotaxis (Xu et al., 2021; Xu et al., 2017; Zhang et al., 2008). It is widely believed that chemotaxis across a broad range of chemoattractant concentrations relies on precise spatiotemporal regulation of Ras activity through coordinated adaptation mechanisms. However, the molecular basis by which GPCR signaling controls the spatial and temporal dynamics of RasGAPs for adaptation during gradient sensing remains poorly understood.

In the present study, we characterized the spatiotemporal dynamics of gradient sensing in the cytoskeleton-free, gradient-sensing cells with or without C2GAP1 (*c2gapA^−^*) in response to steady and/or changing gradients. We found that *c2gapA^−^* cells fail in concentration-dependent adaptation process during gradient sensing. We further revealed an interaction between C2GAP1 and Gα2 and demonstrate their essential role in mediating gradient sensing, enabling rapid orientation in steady gradients, and efficient reorientation in dynamically changing gradients.

## MATERIALS AND METHODS

### Cell lines, cell growth and differentiation

Cell growth and transformation were carried out as previously described (12). Cell expressing the protein of interest was selected by growth in the presence of 20 μg/ml geneticin (Sigma, Steinheim, Germany) or 10 µg/ml blasticidin S, and/or hygromycin (Sigma, Steinheim, Germany) with the requirement of double selection. For differentiation, log-phase vegetative cells were harvested from shaking culture (5×106 cells/ml) and washed twice with developmental buffer (DB: 5 mM Na_2_HPO_4_, 5 mM KH_2_PO_4_, 2 mM MgSO_4_, and 0.2 mM CaCl_2_). Cells were resuspended at 2×10^7^ cells/ml in shaking flask at 100 rpm and allowed to differentiate with 75 nM adenosine 3’:5’-cyclic monophosphate (cAMP) (Sigma Aldrich, St. Louis, MO) pulses at 6 min intervals for 5-7 hours or longer to obtain chemotactic cells. Differentiated cells were diluted to 2 × 10^7^ cells/ml in DB buffer with 2.5 mM caffeine and shaken at 200 rpm for 15 min prior to the experiments.

### Imaging and data processing

Differentiated cells (5 × 10^4^) in DB buffer with 2.5 mM caffeine were plated and allowed to adhere to the cover glass of a 4-well or a 1-well chamber (Nalge Nunc International, Naperville, IL) for 10 min and then covered with DB buffer for the live cell imaging experiment. If necessary, cells were treated with 5.0 μM Lat B (Molecular Probes, Eugene, OR) for 10 min prior to the experiments. Cells were imaged using a Carole Zeiss Laser Scanning Microscope Zen 780 (Carl Zeiss, Thornwood, NY), with a 40x/NA 1.3 Oil DIC Plan-Apochromatic objective. To obtain cytoskeleton-free, immobile cells, cells were treated with 5 μM latrunculin B (Thermo Fisher Scientific Inc., Waltham, Massachusetts) at a final concentration at room temperature for 10 min prior to the experiments. To establish a steady gradient for chemotaxis or gradient-sensing measurements, we set FemtoJet (FemtoJet and micromanipulator 5171, Eppendorf, Germany) with Pc = 70 and Pi = 70 to ensure the injection of a constant and small volume of cAMP and Alexa 594 into a one-well chamber (Thermo Fisher Scientific Inc., Waltham, Massachusetts) as previously described (Xu et al., 2005). Under this condition, a stable gradient was established within 100 μm around the tip of the micropipette. To visualize gradient, cAMP was mixed with Alexa 594 (Thermo Fisher Scientific Inc., Waltham, Massachusetts) at a final concentration of 0.1 mg/ml. To suddenly expose a cell to a stable gradient or reapplication of the identical gradient to the same cell, a micropipette filled with a mixture of cAMP and 0.1 g/ml Alexa 594 linked with a FemtoJet was positioned 1000 μm away from the cells and then was quickly moved to a position within 100 μm to the cells. During the experiments, we only changed the distance between the micropipette and the cells. The speed of movement determines how fast a stable gradient can form around a cell. Images were processed and analyzed by Zen 780 software. Images were further processed in Adobe Photoshop (Adobe Systems, San Jose, CA), and the intensity of the ROI (region of interest) was explored and analyzed with Microsoft Office Excel (Redmond, WA). To measure the membrane translocation of the indicated protein, we first measured the intensity change of the cytoplasm in response to uniformly applied stimuli or in the gradient over time. To obtain the relative intensity change of each individual cell during the time lapse, we divided its intensity at given time (I_t_) by its intensity at time 0 (I_0_); consequently, the relative intensity of any cells at time 0 became 1. To compensate for significant photobleaching that occurs with long-time acquisition, we also normalized the intensity relative to the photobleaching of the cells. We then divided the normalized intensity at time 0 s (I_0_) by the normalized intensity at the given time (I_t_) to convert the normalized intensity change of the cytoplasm to membrane translocation. Lastly, we calculated and presented the mean and standard deviation (Mean ±SD) of peak membrane translocation from more than 5 independent cells.

### Immunoblotting

Mouse monoclonal anti-GFP antibodies were purchased from Clontech Laboratories, Inc. (Mountain View, CA). Anti-Gα2 rabbit antibody from Peter Devreotes, Johns Hopkins University. Anti-pan Ras mouse monoclonal antibody from EMD Millipore (Germany) was used to detect *D. discoideum* Ras proteins. HRP-conjugated anti-mouse or anti-rabbit IgG was obtained from Jackson ImmunoResearch (West Grove, PA).

### Membrane translocation assay

As previously described (Xu et al., 2022a), differentiated C2GAP1-YFP-expressing cells were washed twice with PM buffer (5 mM Na_2_PO4, 5 mM KH_2_PO_4_, and 2 mM MgSO_4_) with 2.5 mM caffeine, resuspended to 2 × 10^7^ cells/ml in PM buffer, and kept on ice prior to stimulation. For the assay, cells were transferred to a medical cup and rotated at 200 rpm at room temperature for 3 min and then stimulated with cAMP at the indicated final concentrations at time 0 s. At each indicated time points, 0.2 ml aliquots were collected and lysed by passing them through a 2-μm pore-size filter unit (Millipore Sigma, Rockville MD) into 1 ml of ice-cold PM buffer. The plasma membrane (PM) fractions were isolated by centrifugation at 15,000 × *g* for 1 min. After carefully removing the supernatants, the PM pellets were solubilized in 50 µl SDS sample buffer (Bio-Rad, Hercules, CA) and subjected to SDS–PAGE and immunoblotting to detect PM-associated C2GAP1–YFP.

### Immunoprecipitation assay

Differentiated C2GAP1-YFP expressing cells were suspended 2 × 10^7^ with PM buffer with 2.5 mM caffeine and kept on ice before assay. Cells were stimulated with 10 μM cAMP. 0.5 ml aliquots of cells at indicated time points were lysed with 10 ml immunoprecipitation buffer (IB, 20 mM Tris, pH8.0, 20 μM MgCl_2_, 10% glycerol, 2 μM Na_3_VO_4_, 0.25 % NP40, and one tablet of Complete 1× EDTA-free proteinase inhibitor) for 30 min on ice. Cell extracts were centrifuged at 16,000 × *g* for 10 min at 4 °C. Supernatant fractions were collected and incubated with 25 μl anti-GFP agarose beads (Proteintec, Rosemont, IL) at 4 °C for 2 hours. Beads were washed four times with immunoprecipitation buffer and proteins were eluted by boiling the beads in 50 μl SDS sample buffer (Bio-Rad, Hercules, CA).

### AlphaFold3-assisted analysis of Gα2-C2GAP1 interaction using NIH Biowulf cluster

Protein–protein complex models were generated using AlphaFold3 (DeepMind, 2024) installed as singularity container with af3 wrapper script (https://github.com/NIH-HPC/af3) on the NIH Biowulf high-performance computing cluster. Model parameters were downloaded from Google DeepMind with permission for non-commercial use. Protein sequences were obtained from dictyBase.org. Jobs were submitted using the default AlphaFold3 (AF3) settings. Predictions were generated for Gα2 (-GDP or -GTP bound), C2GAP1, Gβ, and Gγ along with six interactions, including Gα2GβGγ(-GDP or -GTP bound), C2GAP1-Gα2 (-GDP or -GTP bound), and Gα2GβGγ(-GDP or -GTP bound)-C2GAP1. Five predictions were generated for each job with the top-ranked structure saved in CIF and JSON format and then accessed and visualized in ChimeraX version 1.11.1. Model accuracy was annotated in ChimeraX using the AlphaFold palette, with color-scheme mapped to per-residue confidence scores (pLDDT) accessed from the B-factor column of each respective CIF file. The parameters of the binding affinity for predicted structures were calculated using PRODIGY based on instructions from https://github.com/haddocking/prodigy (Xue et al., 2016).

## RESULTS

### *c2gapA^−^* cells fail to exhibit chemoattractant concentration-dependent adaptation upon exposure to a steady gradient

Chemotaxis is a coordinated process involving three key aspects: gradient sensing machinery, cell polarity, and cell migration. Gradient sensing can be uncoupled from initial cell polarity and cell migration (Parent and Devreotes, 1999). For example, a cell treated with actin polymerization inhibitors loses filamental actin-based cytoskeleton and initial polarity, becomes immobile, yet it retains the ability to sense a chemoattractant gradient. This immobile cell provides a simplified experimental system to specifically investigate the gradient-sensing mechanisms. Using this approach, we previously determined the spatiotemporal dynamics of key signaling components during the polarization processes, including a polarized PIP_3_ production/accumulation in the front of cells in response to exposure to a steady chemoattractant gradient (Xu et al., 2005). A defining feature of gradient-sensing cells is their ability to efficiently establish intracellular polarity, manifested as a sharp accumulation of PIP_3_ in the front of cells toward the source of a chemoattractant, at a concentration-dependent fashion (Meier-Schellersheim et al., 2006; Xu et al., 2005). To understand the role of C2GAP1 in adaptation process of gradient sensing, we first monitored PIP_3_ dynamics using PIP_3_ biosensor, PH_Crac_-GFP, in both WT and *c2gapA^−^* exposed to cAMP gradients of varying concentrations (**Figure 1A** and **Video S1**). WT and *c2gapA^−^* cells expressing PH_Crac_-GFP (green) were treated with 5 μM Latrunculin B for 10 min prior to imaging to diminish cell polarity and cell migration. To visualize cAMP gradients, cAMP was mixed with Alexa594 (red) (Xu et al., 2006). Upon exposure to a relatively low cAMP concentration gradient (100 nM), WT cells exhibited a single-phase PIP_3_ response, characterized by PH_Crac_-GFP translocation only to the front and lateral regions of cells, with minimal accumulation at the rear. The accumulation of PH_Crac_-GFP facing the gradient is also called PH crescent (Parent and Devreotes, 1999). In contrast, exposure to a high, saturating cAMP gradient (10 μM) induced a biphasic response in WT cells, consisting of an initial, uniform PH_Crac_-GFP translocation to the entire cell periphery, followed by withdrawal from the plasma membrane and a second, localized PH_Crac_-GFP accumulation (PH crescent, a PIP_3_ polarization) at the front facing the gradient (**Figure 1A**, top panel). This behavior reflects a typical adaptation process during gradient sensing and is consistent with previous reports (Meier-Schellersheim et al., 2006; Xu et al., 2005). We next examined PIP_3_ dynamics in *c2gapA^−^* cells exposed to either 100 nM or 10 μM cAMP gradients. In contrast to WT cells, *c2gapA^−^* cells displayed persistent PH_Crac_-GFP accumulations at the cell front under both concentrations, indicating a failure in adaptation during gradient sensing (**Figure 1A**, lower panel). The quantitative analysis of PIP_3_ dynamics during gradient sensing, assessed by measuring PH-GFP localization in the front and back (rear) regions of cells in the multiple cells, further confirms this conclusion (**Figure 1B**). The above result demonstrates that C2GAP1 plays an essential role in PIP_3_ adaptation during gradient sensing.

**Figure 1.**
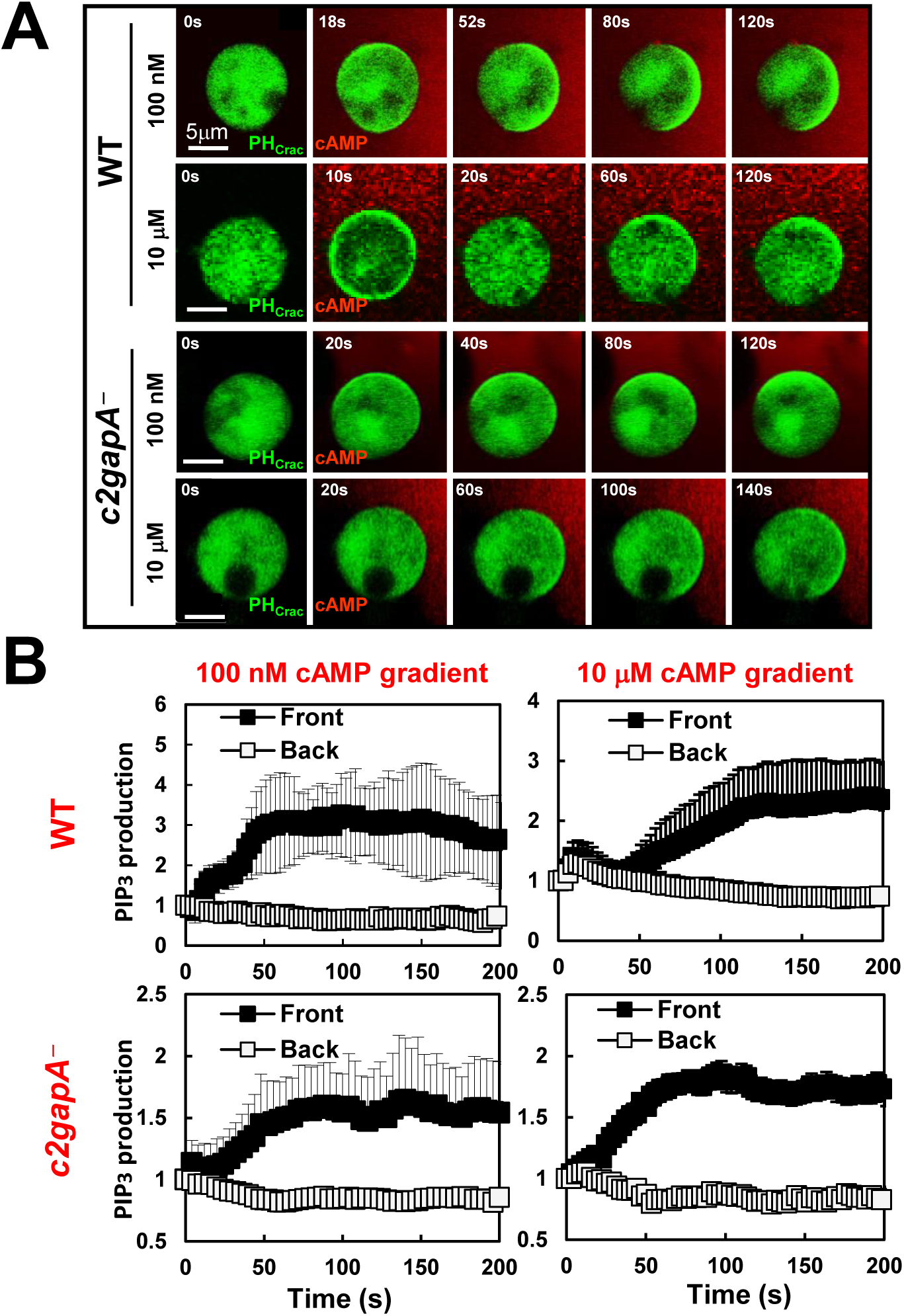
*c2gapA^−^* cells fail to display a concentration-dependent adaptive PIP_3_ dynamics of gradient sensing upon exposure to a steady gradient. **A.** Montage shows PIP_3_ dynamics in wild-type (WT) and *c2gapA^−^* cells upon exposure to steady cAMP gradients generated from the sources of either 10 μM or 100 nM, respectively. Cells expressed PIP_3_ biosensor, PH_Crac_-GFP (green), were treated with 5 μM Latrunculin B for 10 min prior to the experiments. To visualize cAMP gradients, cAMP at the indicated concentrations was mixed with Alexa594 (red). See **Video S1** for complete set of cell responses. Scale bar = 5 μm. **B.** Normalized PIP_3_ production in the front and back of the cells exposed to steady gradients is shown. The PIP_3_ intensity in the front and region at time 0 s was normalized to 1. Mean ± SD is shown. N = 5 and 5 in WT and *c2gapA^−^* cells, respectively, at the indicated concentrations.

### *c2gapA^−^* cells show an altered PTEN dynamics upon exposure to a steady gradient

Phosphatase PTEN is a key molecule that dephosphorylates PIP_3_ and modulates PIP_3_ production during gradient sensing and chemotaxis (Funamoto et al., 2002; Iijima and Devreotes, 2002). We next monitored PTEN dynamics in WT and *c2gapA^−^* cells upon exposure to steady cAMP gradients at varying concentrations (**Figure 2A** and **Video S2**). WT and *c2gapA^−^* cells expressing PTEN-GFP (green) were treated with 5 μM Latrunculin B for 10 min prior to imaging. cAMP was mixed with Alexa594 (red) to visualize the gradient. PTEN-GFP localized at the plasma membrane in both unstimulated, resting WT and *c2gapA^−^* cells, as previously reported (Iijima and Devreotes, 2002). Upon exposure to a relatively low cAMP concentration gradient (100 nM), WT cells exhibited a single-phase PTEN response, characterized by withdrawal of PTEN-GFP from the front and lateral regions of cells (**Figure 2A**, upper panel). In contrast, exposure to a high, saturating cAMP gradient (10 μM) induced a biphasic PTEN response in WT cells, consisting of an initial, uniform withdrawal of PTEN-GFP from the entire cell periphery to the cytoplasm, followed by re-localization of PTEN-GFP from the cytoplasm to the rear of cells. This adaptive redistribution of PTEN restricts PIP3 accumulation to the cell front and is a hallmark of proper gradient sensing (Meier-Schellersheim et al., 2006; Xu et al., 2005). We next examined PTEN-GFP dynamics in *c2gapA^−^* cells exposed to either 100 nM or 10 μM cAMP gradients (**Figure 2A**, lower panel). In contrast WT cells, *c2gapA^−^* cells displayed persistent withdrawal of PTEN-GFP at the cell front under both concentrations. This defective PTEN adaptation is consistent with the spatiotemporal PIP_3_ dynamics, resulting in sustained or excessive PIP_3_ accumulation at the leading edge. The quantitative analysis of PTEN dynamics during gradient sensing, assessed by PM-localization of PTEN-GFP in the front and back regions across multiple cells, further verifies the above conclusion (**Figure 2B**). The PTEN dynamics in WT and *c2gapA^−^* cells are consistent with the PIP_3_ dynamics shown in **Figure 1**.

**Figure 2.**
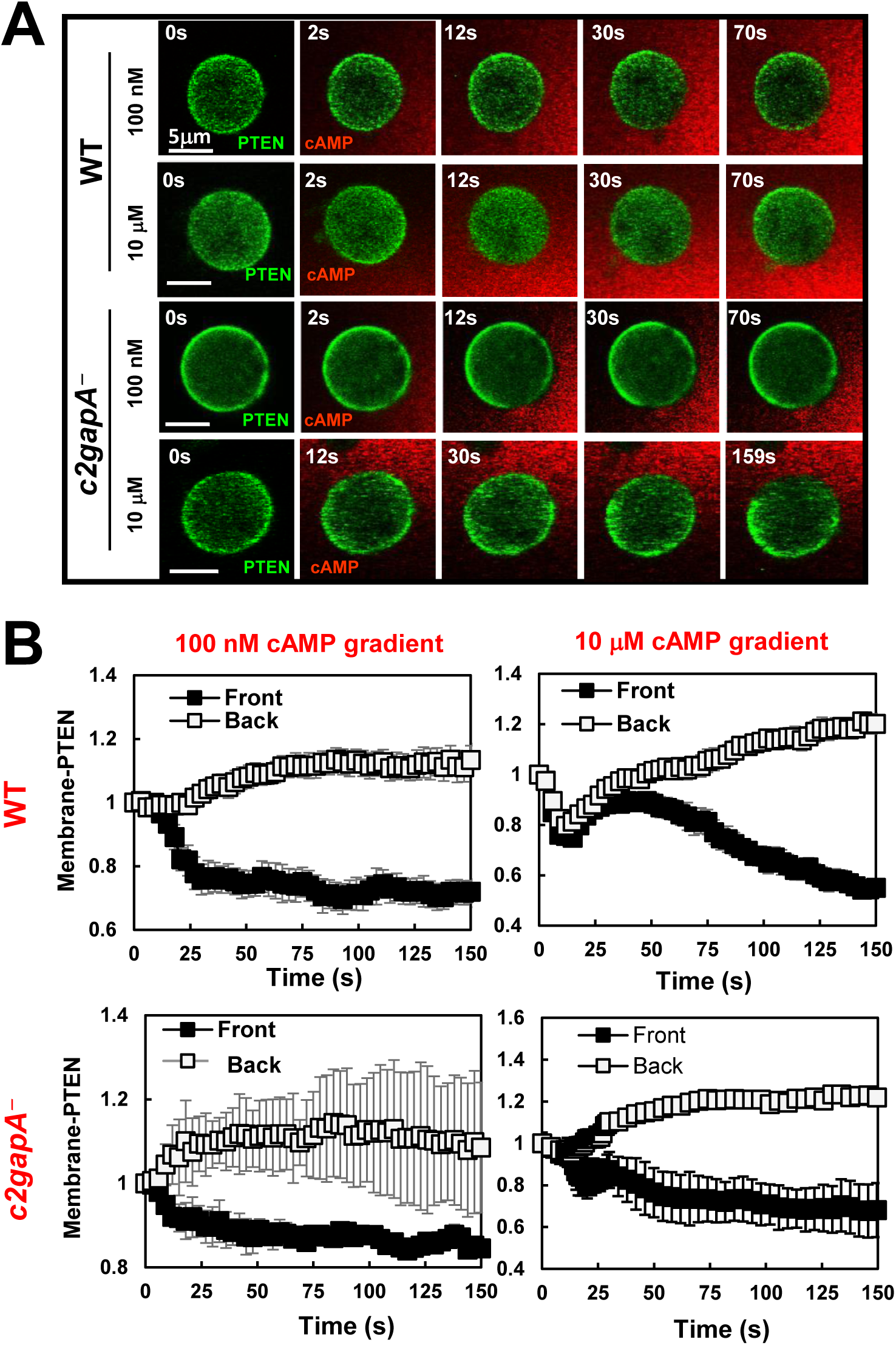
*c2gapA*− cells show altered PTEN dynamics upon exposure to a steady gradient. **A.** Montage shows PTEN dynamics in WT and *c2gapA^−^* cells upon exposure to steady cAMP gradients at the indicated concentrations. Cells expressed PTEN-GFP (green) were treated with 5 μM Latrunculin B for 10 min prior to the experiments. To visualize cAMP gradients, cAMP at the indicated concentrations was mixed with Alexa594 (red). Scale bar = 5 μm. See **Video S2** for complete sets of cell responses. **B.** Normalized intensity of PTEN-GFP in the front and back of the cells upon exposure to steady gradients is shown. The PTEN intensity in the front and region at time 0 was normalized to 1. Mean ± SD is shown. N = 5 and 5 in WT and *c2gapA^−^* cells, respectively, at both indicated concentrations.

### F-actin independent, cAMP concentration-dependent PM targeting of C2GAP1 during gradient sensing

To understand the F-actin-independent, chemoattractant concentration-dependent adaptation in gradient sensing, we next examined whether C2GAP1 translocates to the plasma membrane (PM) in an F-actin independent, cAMP concentration-dependent manner. We first assessed PM translocation of C2GAP1–YFP biochemically in a large population of Latrunculin B–treated cells stimulated with either high (10 μM) or low (10 nM) cAMP. Plasma membrane fractions were isolated before and after cAMP stimulation as previously described (Xu et al., 2022a). A clear cAMP concentration-dependent PM translation of C2GAP1-YFP was observed (**Figure 3A**). Uniform stimulation at a high, saturating concentration of cAMP (10 μM) induced a biphasic PM translocation of C2GAP1: a rapid initial recruitment peaking at ∼10–15 s, followed by dissociation from the PM by ∼60 s, and subsequently a second, sustained PM localization lasting up to 180 s. In contrast, stimulation at a low concentration of cAMP (10 nM) elicited a single-phase response characterized by a slow but prolonged PM recruitment between ∼10 and 30 s, followed by a return to pre-stimulation levels. Quantitative analysis of three independent experiments confirmed these observations (**Figure 3B** and **Figure S1**).

**Figure 3.**
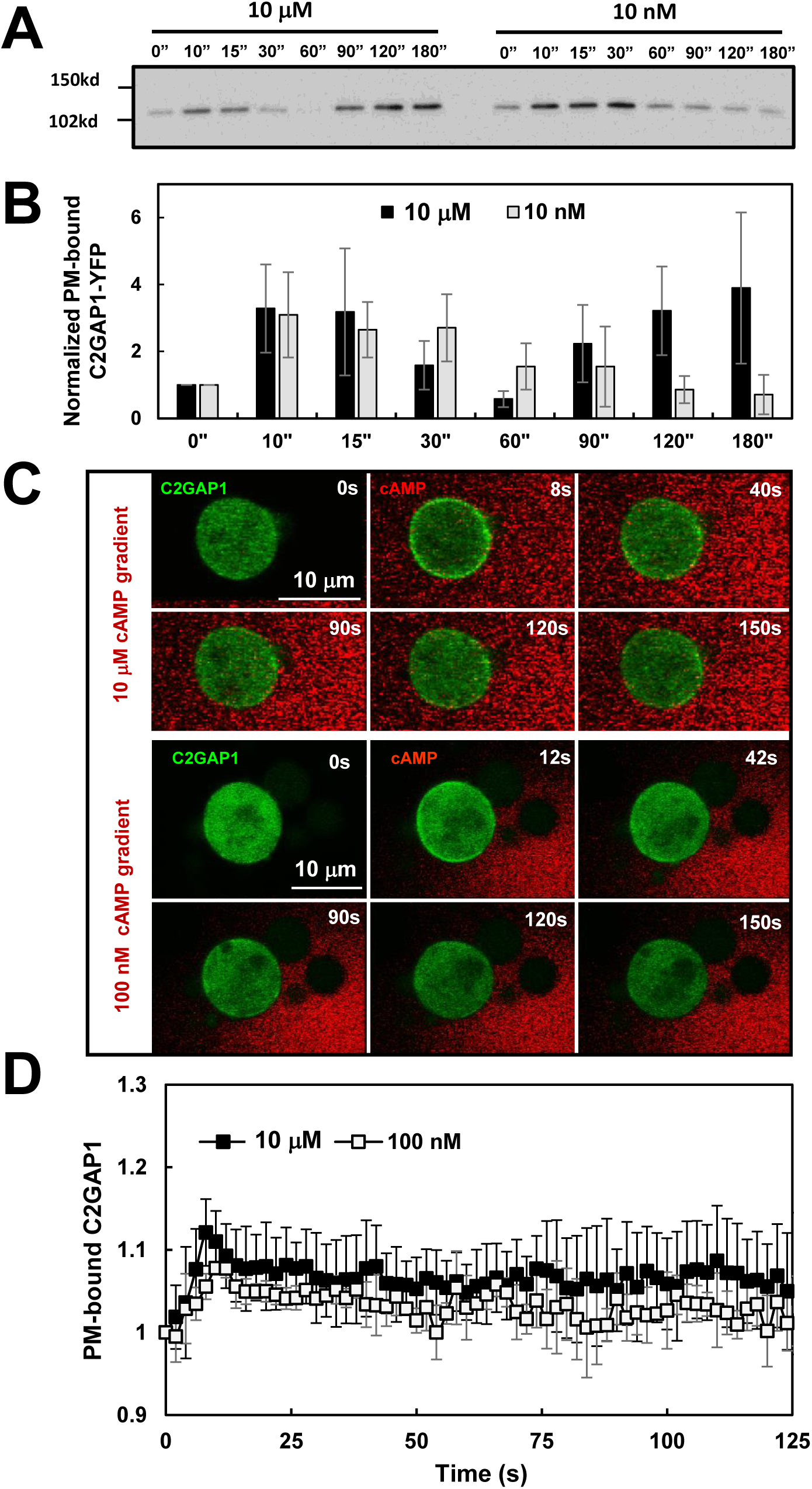
F-actin-independent, cAMP concentration-dependent PM targeting C2GAP1 during gradient sensing. **A.** F-actin-independent, cAMP concentration-dependent plasma membrane translocation dynamics of C2GAP1 upon cAMP stimulation. Cells treated with 5 μM Latrunculin B 10 prior to the experiment and were stimulated with cAMP at the indicated concentration at 0 s. Plasma membrane fractions were collected at the indicated time points and subjected to Western blot analysis to detect C2GAP1-YFP using anti-GFP antibodies. **B.** Normalized plasma membrane (PM) translocation of C2GAP1 upon cAMP stimulation from A and two additional independent experiments (**Figure S1**). C2GAP1 membrane localization at 0 s was normalized to 1. **C.** Montage shows C2GAP1-GFP dynamics in WT cells upon exposure to steady cAMP gradients at the indicated concentrations. Cells expressed C2GAP1-GFP (green) were treated with 5 μM Latrunculin B for 10 min prior to the experiments. To visualize cAMP gradients, cAMP at the indicated concentrations was mixed with Alexa594 (red). See **Video S3** for complete sets of cell responses. **D.** Normalized PM translocation of C2GAP1-GFP upon exposure to steady gradients is shown. The cytosolic intensity of C2GAP1 at time 0 s was normalized to 1. Mean ± SD is shown. N = 5 and 5 in WT cells exposed to cAMP gradients at either 10 μM or 100 nM, respectively.

We next monitored spatiotemporal PM translocation of C2GAP1-YFP in the cell response to exposure to a steady cAMP gradient by confocal microscopy (**Figure 3C** and **Video S3**). *c2gapA^−^* cells expressing C2GAP1-YFP (green) were treated with 5 μM Latrunculin B for 10 min prior to imaging. cAMP was mixed with Alexa594 (red) to visualize the gradient. Upon exposure to a high, saturating cAMP (10 μM) gradient, cells exhibited an initial PM translocation peak around ∼ 8 s, followed by a withdrawal from the PM and maintained a sustained level on the PM (**Figure 3C**, upper panel). Notably, C2GAP1-YFP accumulated more prominently in the front than at the rear. Upon exposure to a low-concentration cAMP (100 nM) gradient, a similar response pattern, although the peak translocation occurs slower at 12 s (**Figure 3C**, lower panel). Quantitative analysis of membrane translocation, assessed by cytosolic C2GAP1 depletion, confirmed these observations and further revealed a concentration-dependent response (**Figure 3D**). Collectively, these results demonstrate that C2GAP1 accumulates in the plasma membrane in an F-actin independent, cAMP concentration-dependent manner during gradient sensing.

### cAMP concentration-dependent inhibitory process of gradient-sensing cells

One key feature of F-actin free, PIP_3_-polarized gradient-sensing cells is that cells accumulate a stronger inhibition in the front of cells facing a steady, persistent gradient (Meier-Schellersheim et al., 2006; Xu et al., 2007). This stronger inhibition was experimentally verified (Xu et al., 2007). The experimental design is shown in **Figure 4A**. Cells initially exposed to a steady gradient displayed stable PH_Crac_-GFP crescents oriented toward the gradient source. Next, the gradient was removed, the PH_Crac_-GFP crescents gradually diminished and disappeared in the front. Upon reapplication of second stimulation (either a uniform or a gradient), PH_Crac_-GFP translocates exclusively at the original rear of the cells. We refer to this behavior as an inversed response. The inverse response was also elicited upon reapplication of the same gradient, whereby cells effectively experienced a higher stimulus at the front relative to the rear. The eventual disappearance of the inversed response, or its reorientation and re-accumulation of PH_Crac_-GFP at the front facing the gradient source, demonstrates that cells retain their ability to sense and respond to the gradient (Xu et al., 2007). We designated this increased sensitivity at the rear of PIP_3_-polarized cells as inverse sensitivity, reflecting the preferential responsiveness of the original back. To further elucidate the nature of inverse sensitivity, we investigated whether it depends on the cAMP concentration of the gradient. To address this, we investigated the inversed sensitivity of PIP_3_-polarized cells exposed to either high (10 μM) or low (100 nM) cAMP gradient. After cells established a stable PH_Crac_ crescent in the gradient, the initial gradient was removed to allow dissipation of front-facing PH_Crac_-GFP accumulation. An identical, second gradient was reintroduced, and cellular response was monitored (**Figure 4B**, **Video S4**). We found that PIP_3_-polarized cells exposed to 10 μM cAMP gradients frequently (∼ 93.6%) displayed inverse sensitivity (**Figure 4B**, left upper panel). In contrast, only ∼8.2% of cells exposed to 100 nM cAMP gradients exhibited an inverse response (**Figure 4B**, left lower panel). The selected areas shown in the montages indicate the regions used for quantitative measurement of cAMP gradient sensing and PH_Crac_-GFP PM translocation. Quantitative analysis confirmed that identical gradients were applied to the cells and revealed the temporal and spatial dynamics of PIP_3_ accumulation at both the front and back of the cells (**Figure 4C**). Taken together, these results demonstrate that an F-actin-independent, cAMP concentration–dependent inhibitory process is an intrinsic component of gradient sensing.

**Figure 4.**
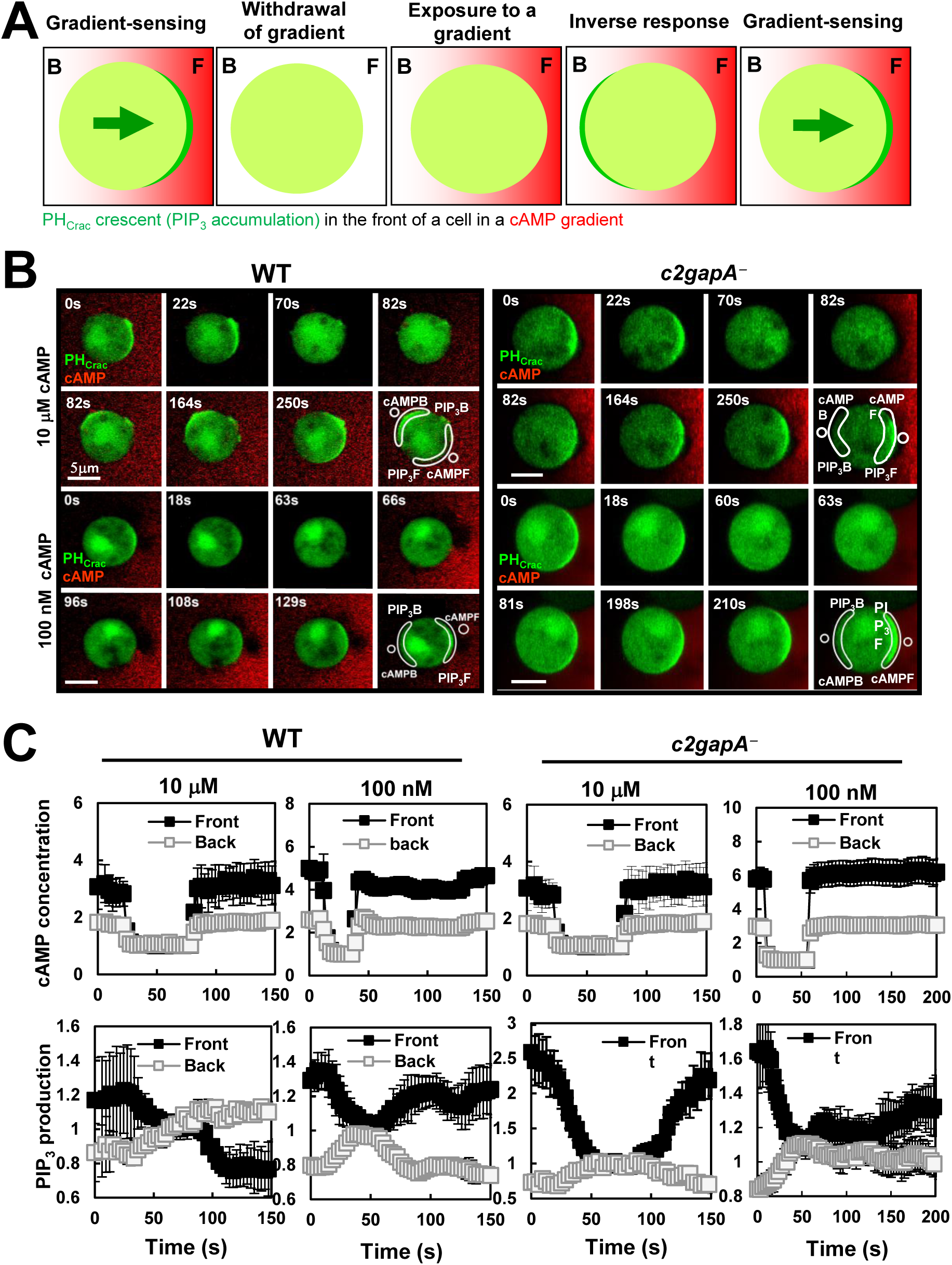
PIP_3_-polarized *c2gapA^−^* cells fail to display a concentration-dependent, higher inhibition in the front of gradient-sensing WT cells. **A.** Montages show PIP_3_ dynamics in PIP_3_-polarizaed WT and *c2gapA*^-^ cells upon removal of the original gradient and then re-exposure to a second, identical gradient at the indicated concentrations. Cells expressing PIP_3_ probe, PH_Crac_-GFP (green), were treated with 5 μM Latrunculin B for 10 min prior to the experiments. To visualize cAMP gradient, cAMP at the indicated concentrations was mixed with Alexa594 (Red). Gradient sensing capability was indicated by the accumulation of PH-GFP in the front of the cells facing the source of a cAMP gradient. See **Video S4** for complete sets of cell responses. **B.** Normalized intensity of PIP_3_ and cAMP concentrations the front and back of c*2gapA^-^*cells upon removal and re-exposure to identical gradients. cAMP concentrations and PIP_3_ intensity at the time of completely withdrawn from plasma membrane upon the removal of the initial gradient was normalized to 1. Mean ± SD is shown. N = 5 and 9 in the gradients of either 10 μM or 100 nM cAMP, respectively.

### *c2gapA^−^* cell fails to display a higher inhibition in the front during gradient sensing

We previously showed that C2GAP1 accumulates at the plasma membrane in a cAMP concentration–dependent manner in cells exposed to a cAMP gradient (**Figure 3**). We next investigated whether C2GAP1 contributes to the cAMP concentration–dependent inhibitory mechanism. To address this, we examined whether gradient-sensing, PIP_3_-polarized *c2gapA⁻* cells exhibit concentration-dependent inverse sensitivity. *c2gapA^−^* cells expressing PIP_3_ biosensor PH_Crac_-GFP (green) were treated with 5 μM Latrunculin B for 10 min prior to the experiment and cAMP was mixed with Alexa594 (red). Upon exposure to either 10 μM or 100 nM cAMP gradients, *c2gapA⁻*cells displayed PH-GFP accumulation (PH crescents) at the front facing the gradient (**Figure 4A**, right panel; **Video S4**). Removal of either gradient caused PH crescents to gradually diminish and disappear. Upon re-exposure to the identical gradient, PH-GFP translocated to the original front, instead of back, of *c2gapA⁻*cells. The selected areas shown in the montages indicate the regions used for quantitative measurement of cAMP gradient sensing and PIP_3_ accumulation. Quantitative analysis confirmed that identical gradients were applied and revealed the temporal and spatial dynamics of PIP_3_ accumulation at both the front and back of the cells (**Figure 4C**). Collectively, these results demonstrate that C2GAP1 plays an essential role in establishing cAMP concentration–dependent inhibition during gradient sensing.

### Dynamics of C2GAP1 PM localization in response to removal and second application of cAMP gradient

To investigate the mechanism by which C2GAP1 mediates this concentration-dependent inhibition, we monitored plasma membrane localization of C2GAP1 following gradient removal and reapplication (**Figure 5A**; **Video S5**). Chemotactic C2GAP1-YFP (green)–expressing *c2gapA⁻* cells treated with latrunculin B (5 μM) were exposed to a 10 μM cAMP gradient (red). After exposure to a steady gradient for more than 200 s, the gradient was removed and this time point was designated as 0 s. At 54 s, an identical gradient was reintroduced, inducing a second translocation of C2GAP1-YFP to the plasma membrane followed by partial withdrawal.

**Figure 5.**
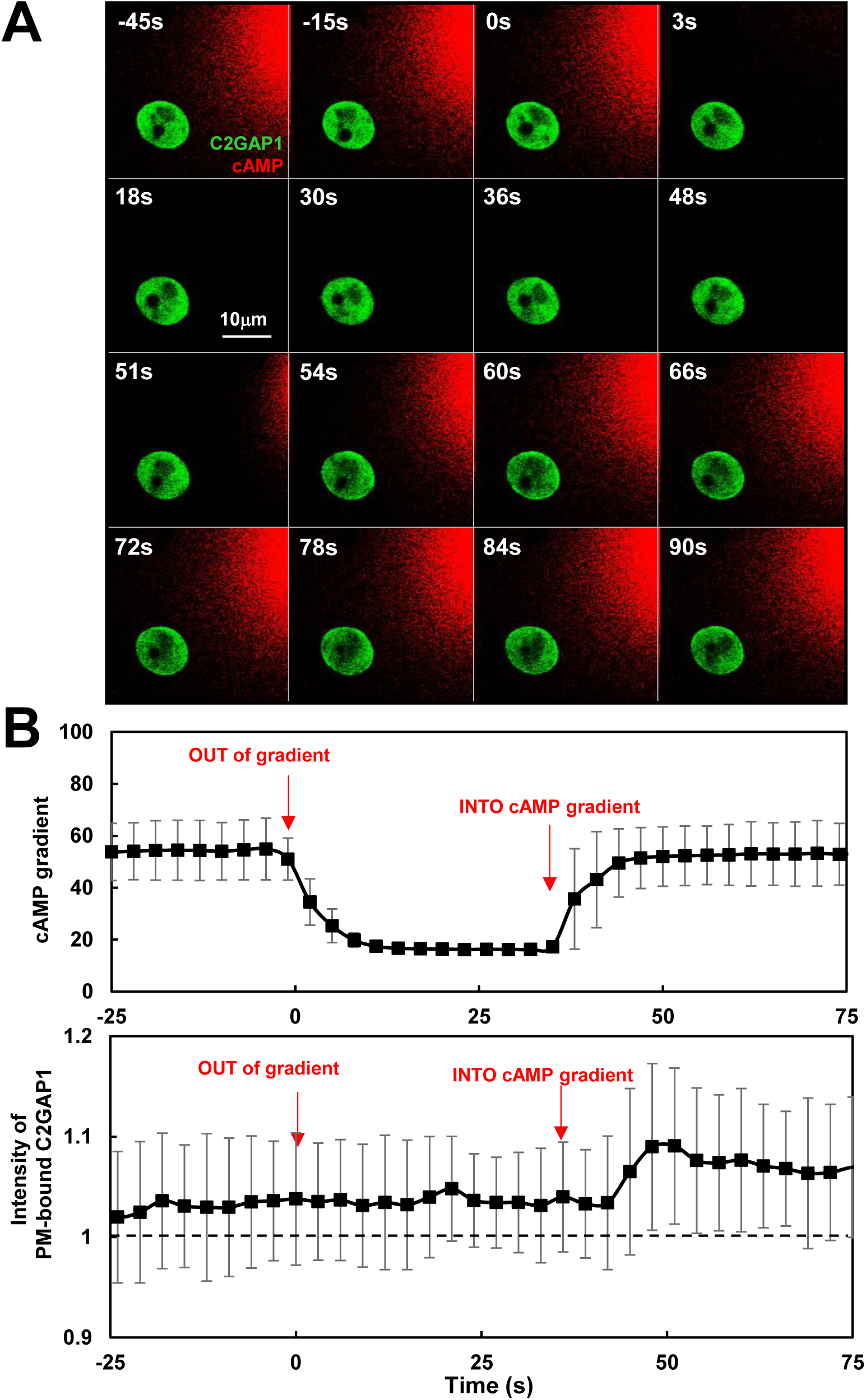
C2GAP1 dynamics in response to removal and second application of cAMP gradient. **A.** Montages show C2GAP1-YFP (green) dynamics in PIP_3_-polarizaed WT and *c2gapA*^-^cells upon removal of the original gradient and then re-exposure to a second, identical gradient at the indicated concentrations. Cells expressing C2GAP1-YFP (green), were treated with 5 μM Latrunculin B for 10 min prior to the experiments. To visualize cAMP gradient, 10 μM cAMP was mixed with Alexa594 (red). See **Video S5** for complete set of cell responses, including exposure to the initial gradient and then removal and reapplication of a second identical gradient. Time 0 s was set at the time last scan before the removal of the existing, original gradient and reapplication of the gradient at time 51s. **B.** Normalized intensity of PIP_3_ and cAMP concentrations the front and back of c*2gapA^-^*cells upon removal and re-exposure to identical gradients. cAMP concentrations and PIP_3_ intensity at the time of completely withdrawn from plasma membrane upon the removal of the initial gradient was normalized to 1. Mean ± SD is shown. N = 5 and 9 in the gradients of either 10 μM or 100 nM cAMP, respectively.

Quantitative analysis confirmed removal of the initial gradient and subsequent reapplication of an identical gradient (**Figure 5B**). Plasma membrane localization of C2GAP1 at the time of initial gradient exposure was normalized to 1; notably, membrane-bound C2GAP1 remained above this level at 0 s, indicating persistent plasma membrane association, consistent with **Figure 3D**. A complete dataset of gradient measurements and C2GAP1 PM translocation during initial exposure, gradient removal, and re-exposure is shown in **Figure S2**. Importantly, C2GAP1-YFP did not significantly dissociate from the plasma membrane following gradient removal, suggesting that C2GAP1 persists at the membrane to provide sustained inhibitory signaling during subsequent gradient sensing.

### C2GAP1 interacts with Gα2 and decreases the activation of heterotrimeric G protein upon cAMP stimulation

To better understand the molecular mechanisms by which C2GAP1 modulates Ras signaling to establish a concentration-dependent inhibitory process in achieving the proper PIP_3_ dynamics, we sought to identify additional molecules that interact with C2GAP1 under different conditions by co-immunoprecipitation (co-IP) assays (**Figure 6A**). Cells expressing YFP protein alone were used as a negative control. cAMP-chemotactic, C2GAP1-YFP expressing cells were treated with either cAMP, hydrolysis-resistant GTPγS, or latrunculin B and then subjected to co-IP experiment and immunoblotting of the indicated proteins. Notably, we identified Gα2 as an interacting partner of C2GAP1 with or without cAMP stimulation (**Figure 6A**). cAMP chemoattractant-competent cells express endogenous level of Gα2 that is a subunit of heterotrimeric G protein and essential for gradient sensing in F-actin-free cells (Xu et al., 2005; Xu et al., 2007). Importantly, the association between Gα2 and C2GAP1 persisted in the cells treated with actin polymerization inhibitor, latrunculin B, when F-actin-mediated positive or negative feedback loop was diminished, indicating that that Gα2-C2GAP1 interaction is a core component of the gradient-sensing machinery.

**Figure 6.**
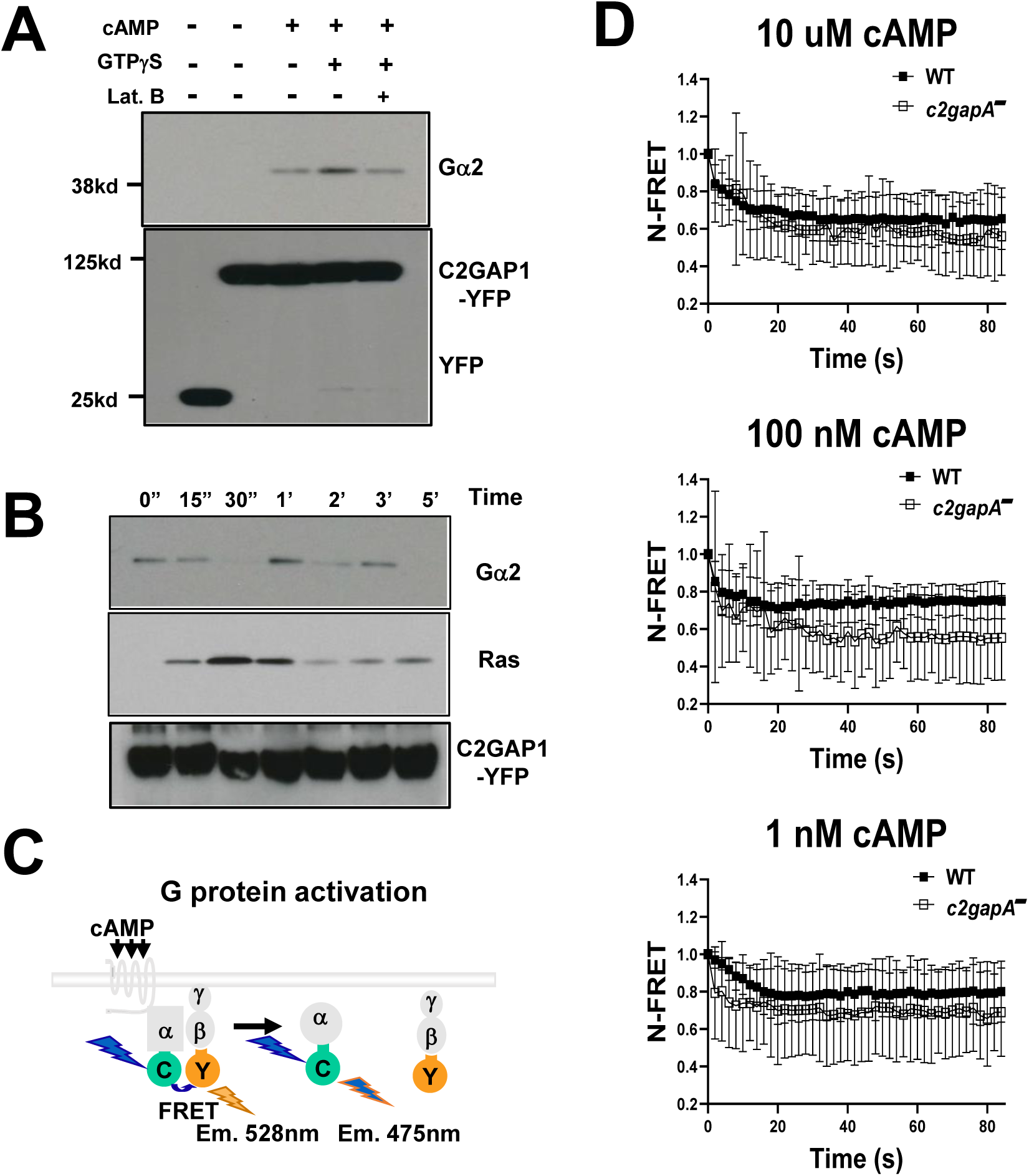
C2GAP1 interacts with Gα2 and decreases the activation of heterotrimeric G protein upon cAMP stimulation. **A.** Interaction of C2GAP1 and Gα2 under various conditions determined by co-immunoprecipitation (co-IP). cAMP-chemotactic competent cells with endogenous level Gα2 and the expression of either YFP (a negative control) or C2GAP1-YFP were wither the treatment of either 10 μM cAMP, 1 mM GTPγS, and/or 5 μM latrunculin B (Lat. B) were lysed and incubated with anti-GFP conjugated agarose beads and subjected to co-IP experiment and western-blotting detection of the indicated molecules. **B.** Dynamic interaction between C2GAP1 and Gα2/Ras upon cAMP stimulation determined by co-immunoprecipitation (co-IP). cAMP-chemotactic competent cells with endogenous level Gα2 and the expression of C2GAP1-YFP were stimulated with cAMP at a final concentration of 10 μM. Aliquots of cells at the indicated time points were subject to co-IP experiment and western-blotting detection of the indicated molecules. C2GAP1-Ras interaction was used to as a positive control. **C.** Schematic illustration of heterotrimeric G protein activation upon cAMP stimulation, measured as a loss of FRET between Gα2-CFP and Gβ-YFP. G protein activation leads to dissociation of Gα2 from Gβγ, resulting in reduced FRET efficiency. **D.** Time course of FRET efficiency changes in WT and *c2gapA⁻* cells following uniform cAMP stimulation. FRET efficiency was quantified using sensitized emission–based FRET analysis.

To further characterize the dynamics of the association, we determined the temporal profile of C2GAP1-Gα2 interaction in response to cAMP stimulation (**Figure 6B**). The C2GAP1−Ras interaction was used as a positive control, and as previously reported, a transient association between C2GAP1 and Ras was observed (Xu et al., 2017). Consistent with the result shown in **Figure 6A**, an association between Gα2 and C2GAP1 in the cells prior to stimulation. This interaction persisted but decreased markedly at 15 s after cAMP stimulation and became undetectable at 30 s, followed by a second peak at approximately 1 min post stimulation and subsequent oscillatory behavior. These interaction dynamics are consistent with previous observations that a fraction of C2GAP1 localizes to the plasma membrane and protrusive regions in resting cells, and that cAMP stimulation induces an initial plasma membrane translocation of C2GAP1 at ∼15 s, followed by withdrawal at ∼30 s, a second translocation at ∼1 min, and subsequent oscillatory recruitment (Xu et al., 2021; Xu et al., 2017). Notably, the second peak of the Gα2–C2GAP1 interaction coincided with the second wave of C2GAP1 plasma membrane localization during gradient sensing.

To assess the role of Gα2−C2GAP1 interaction in heterotrimeric G protein activation, we measured G protein activation by monitoring fluorescence resonance energy transfer (FRET) between Gα2-CFP (FRET donor) and Gβ-YFP (FRET acceptor) in WT and *c2gapA^−^* cells by confocal live cell imaging previously reported (Xu et al., 2005). A schematic illustrating G protein activation upon cAMP stimulation, measured as a loss of FRET-between Gα2-CFP and Gβ-YFP, is shown in **Figure 6C**. Sensitized emission of FRET efficiency was used to quantify FRET loss (Xu et al., 2016). In WT cells, cAMP stimulation at a final concentration of 10 μM induced a decrease in FRET efficiency (**Figure 6D**), consistent with previous reports (Xu et al., 2025; Xu et al., 2016). *c2gapA⁻* cells exhibited a notable greater loss of FRET in response to the same 10 μM cAMP stimulation. This enhanced FRET loss was also observed in *c2gapA⁻* cells stimulated with lower concentrations of cAMP (100 nM and 1 nM). Together, these results demonstrate that the Gα2–C2GAP1 interaction sustains C2GAP1 at the plasma membrane to inhibit Ras signaling for proper adaptation, while simultaneously attenuating heterotrimeric G protein activation in response to cAMP stimulation.

### Simulation of Gα2-C2GAP1 interaction

In resting cells, the majority of heteromeric G protein exists in the inactive GDP-bound form, Gα2(GDP)Gβγ. Saturating cAMP stimulation (10 μM cAMP; F**igure 6B**) induces maximal dissociation of the heterotrimer, leading to activation of Gα2-GTP and release of free Gβγ. Notably, Gα2–C2GAP1 association is detected both before and after cAMP stimulation, indicating that C2GAP1 can interact with Gα2 in either its inactive (GDP-bound) or active (GTP-bound) state. To gain mechanistic insight into this dynamic interaction, we modeled the structures and binding properties of Gα2 and C2GAP1 using AlphaFold 3 (**Figure 7**). We first predicted the structures of Gα2 in its GDP- and GTP-bound forms (Figure 7A). Gα2 adopts the canonical architecture composed of a Ras-like GTPase domain and an α-helical domain, separated by a deep cleft that accommodates GDP or GTP. The switch regions of Gα2 are similarly conserved (**Figure 7B**) (Van Eps et al., 2006). These predicted structures show substantial overlap with mammalian Gαi2 (**Figure S3A**). We also modeled the Gβ and Gγ subunits: Gβ forms a seven-bladed β-propeller, with each blade consisting of four-stranded antiparallel β-sheets, whereas Gγ adopts a coiled-coil structure (**Figure S3B**), both closely resembling their mammalian counterparts (**Figure S3C**). The predicted heterotrimeric G protein complex (Gα2Gβγ) exhibits a conserved overall architecture relative to mammalian structures (Figure S3D) (Lambright et al., 1996).

**Figure 7.**
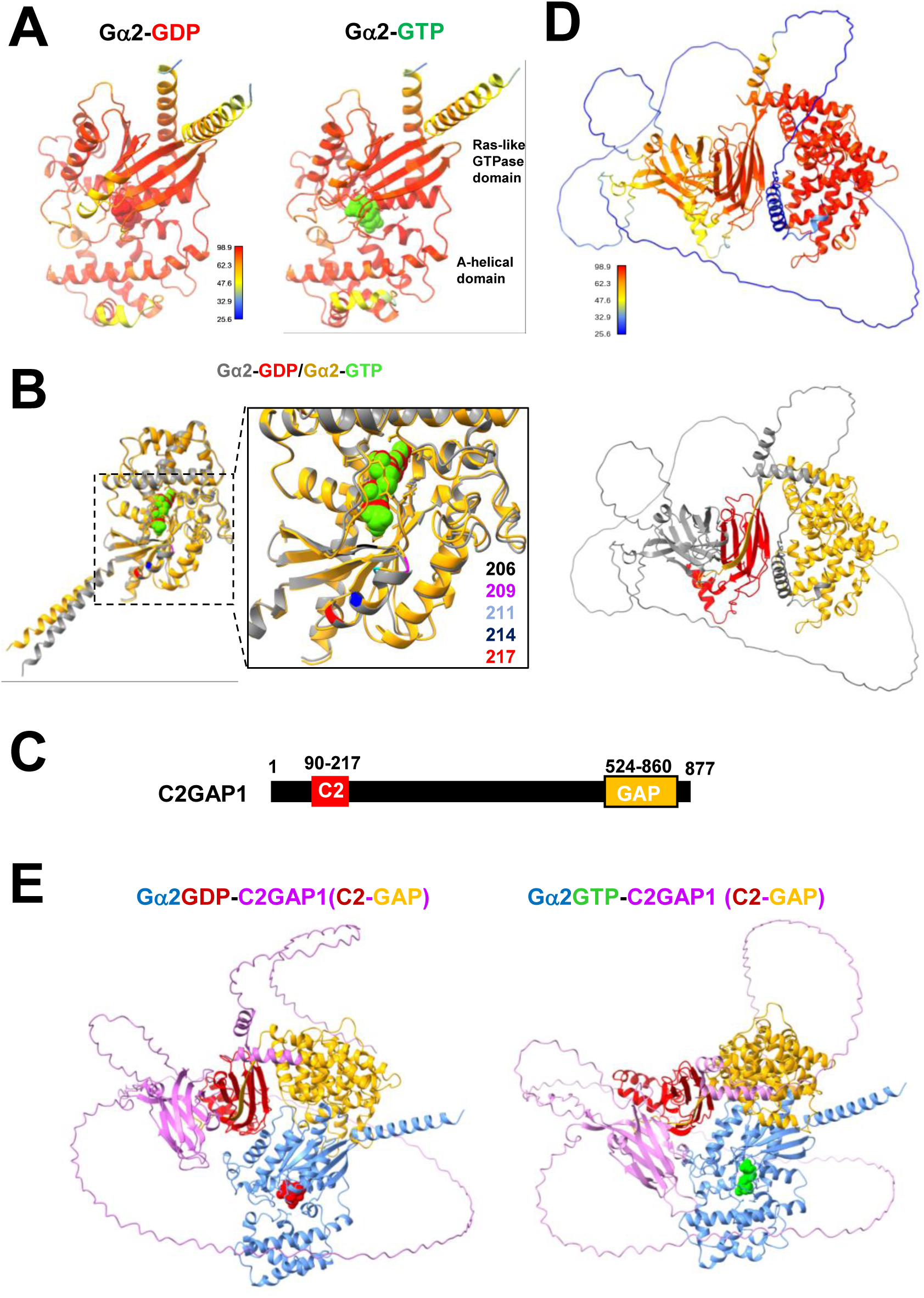
Simulation of the interaction between Gα2 and C2GAP1. **A.** Predicted structures of the heterotrimeric Gα2 subunit in complex with GDP (red spheres) or GTP (green spheres). Gα2 is composed of a Ras-like GTPase domain and an α-helical domain, separated by a deep cleft that accommodates GDP or GTP. **B.** Alignment of Gα2-GDP (grey) and Gα2-GTP (gold). The enlarged area highlights key residues in the switch region, with residue numbers displayed in matching colors. **C.** Domain architecture of C2GAP1, comprising a C2 domain and a GAP domain. **D.** Predicted structures of C2GAP1, with the prediction confidence scale shown in the upper panel and domain composition in the lower panel; the C2 domain is shown in red and the GAP domain in gold. **E.** Predicted Gα2–C2GAP1 complexes, showing Gα2 (blue) in GDP-bound (red) or GTP-bound (green) states and the C2 (red) and GAP (gold) domains of C2GAP1 (magenta).

With these conserved structural features established for *Dictyostelium* heterotrimeric G proteins, we next predicted the structure of C2GAP1 (**Figure 7C–D**). The domain architecture for both the C2 and RasGAP domains were predicted with high confidence. We then modeled the complexes between C2GAP1 and Gα2 in either the GDP- or GTP-bound state (**Figure 7E**). In both nucleotide states, the C2 and GAP domains interact with the Ras GTPase domain of Gα2. Notably, an additional region of C2GAP1 located between the C2 and GAP domains provides extra contacts with the GTPase domain of Gα2 specifically in the GTP-bound state (**Figure S4A**). Binding free-energy calculations yielded predicted values of −9.8 kcal/mol for C2GAP1–Gα2-GDP and −11.2 kcal/mol for C2GAP1–Gα2-GTP, corresponding to inferred dissociation constants of ∼6.8 × 10⁻⁸ M and ∼6.3 × 10⁻⁹ M, respectively (**Figure S4B**). By nature of the simulations, these values should be interpreted qualitatively rather than quantitatively. Nevertheless, the results indicate an approximately one-order-of-magnitude stronger affinity of C2GAP1 for Gα2-GTP compared with Gα2-GDP. Together, these simulations support a model in which C2GAP1 associates with Gα2 in both inactive and active states but preferentially binds activated Gα2-GTP. This enhanced interaction provides a molecular mechanism for the increased plasma membrane recruitment of C2GAP1 during cAMP gradient sensing.

### *c2gapA^−^* cells display significantly impaired reorientation in response to a changing gradient

In the natural environment, chemoattractant gradients are often dynamic, and efficient reorientation is a key aspect of gradient sensing and subsequent chemotaxis. To investigate this, we designed an experiment to quantitatively measure the reorientation dynamics of both WT and *c2gapA⁻* cells in response to a changing gradient. Cells were first allowed to chemotax in a steady gradient for several minutes, after which the gradient was reversed, as illustrated in **Figure 8A** (see **Video S7** for complete sets of cell responses). We observed four distinct cell behaviors: (1) continuous movement in the same direction (SD); (2) formation of a new pseudopod at the original trailing edge and migration toward the new gradient direction (NP); (3) turning of 180° and migrating toward the new gradient direction (TN); and (4) no movement (NM). We then quantitatively measured the time required for each of these behaviors in response to gradient changes at three different cAMP concentrations (10 μM, 100 nM, and 1 nM; **Figure 8B**). In 10 μM and 100 nM gradients, *c2gapA⁻* cells required significantly longer times to adjust to the new gradients compared with WT cells. This difference diminished at lower concentrations. At 1 nM, no significant difference was observed between WT and *c2gapA⁻* cells for SD, NP, or NM behaviors; however, *c2gapA⁻*cells displayed significantly faster turning and migration toward the new gradient direction. These results are consistent with previous reports showing that *c2gapA⁻* cells exhibit concentration-dependent chemotaxis: impaired chemotaxis in high, saturating gradients, similar performance at medium concentrations, and enhanced chemotaxis at low or sub-sensitive concentrations (Xu et al., 2021; Xu et al., 2017). In conclusion, our data indicates that C2GAP1 is critical for rapid reorientation, particularly in gradients at medium to high concentrations.

**Figure 8.**
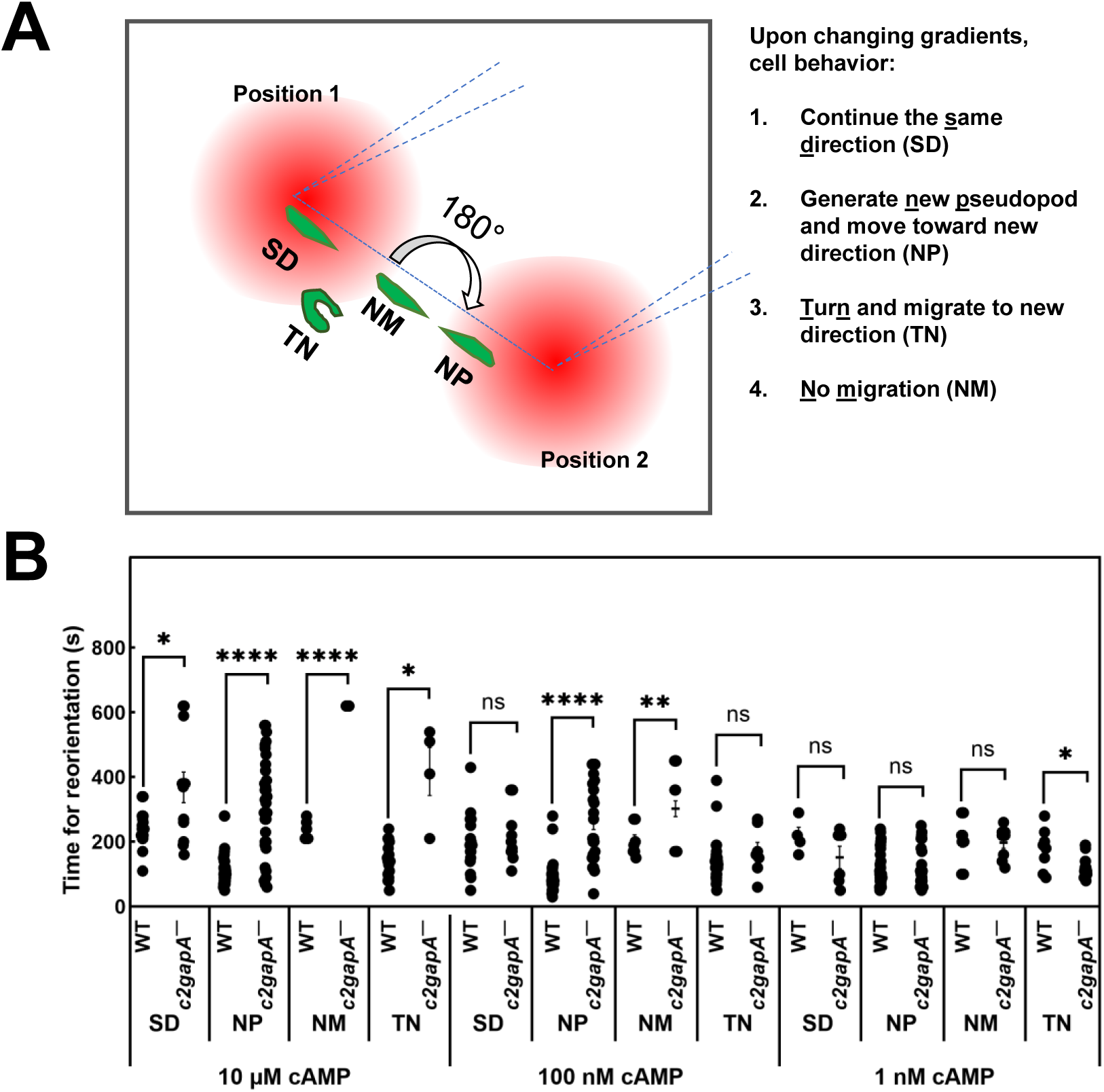
*c2gapA^−^* cell displays significant impaired reorientation in response to a changing gradient. **A.** Schematic of the changing-direction experiment used to assess responses of WT and *c2gapA⁻* cells (see **Video S7** for complete cell responses). Four distinct behaviors were observed: (1) continuous migration in the same direction (SD); (2) formation of a new pseudopod at the original trailing edge followed by migration toward the new gradient direction (NP); (3) turning of 180° and migrating toward the new gradient direction (TN); and (4) no movement (NM). **B.** Time (s) required for reorientation in WT and *c2gapA^−^* cells upon the changing direction of the gradient.

## DISCUSSION

Eukaryotic cells sense and migrate through chemoattractant gradients with enormous concentration range, such as 10^-5^ to 10^-9^ M cAMP in *D. discoideum* and 10^-5^ to 10^-9^ M SDF1a or fMLP in neutrophils. To migrate effectively across such broad concentration ranges, cells employ adaptation mechanisms, by which cells adapt to current stimulus while maintaining sensitivity to stronger signals, enabling continuous movement up a gradient. Chemotaxis involves three conceptually distinct yet interconnected processes: gradient sensing, cell polarity, and cell migration. Among these, gradient sensing provides the foundation for directional migration. Although many components acting through the F-actin–based cytoskeleton have been shown to play pivotal roles in chemotaxis, the core elements of the gradient-sensing machinery and the molecular mechanisms underlying adaptation remain incompletely understood. In this study, we identify a Gα2–C2GAP1 interaction and demonstrate its essential role in mediating adaptation during gradient sensing and in promoting efficient orientation in dynamic gradients.

The essence of gradient sensing is the ability to detect an extracellular gradient and establish an intracellular polarized response. Latrunculin B–treated, cytoskeleton-free, immobile *Dictyostelium* cells retain the capacity to sense gradients and therefore provide a simplified system for specifically investigating gradient sensing (Parent and Devreotes, 1999). Using these immobile cells, the spatiotemporal dynamics of GPCR cAR1–mediated sequential signaling events—including extracellular gradient strength, heterotrimeric G protein activation, Ras activation, and PIP_3_ production—have been monitored upon exposure to a steady gradient (Kortholt et al., 2013; Meier-Schellersheim et al., 2006; Xu et al., 2005; Xu et al., 2007) (**Figure S5**). Upon gradient exposure, cells experience higher chemoattractant concentrations at the front than at the back, triggering rapid and sustained G protein activation that is stronger at the front (Meier-Schellersheim et al., 2006; Xu et al., 2005). This observation indicates that adaptation occurs downstream of G protein activation and that G protein activation reflects the local chemoattractant concentration.

Upon exposure to a steady gradient, cells establish intracellular polarity, manifested by a sharp accumulation of PIP_3_ at the front of the cell facing the chemoattractant source in a concentration-dependent manner. When exposed to relatively low cAMP gradients (<100 nM), cells exhibit a single-phase PIP_3_ production and accumulation at the front and lateral regions, with minimal accumulation at the rear. In contrast, exposure to high, saturating cAMP gradients (>1 μM) elicits a biphasic PIP3 response, consisting of an initial uniform PIP_3_ production around the entire cell periphery, followed by its reduction from the plasma membrane and a second phase of PIP_3_ accumulation at the front—representing a typical adaptation process followed by a polarized response. Nevertheless, the polarized PIP_3_ response during gradient sensing is concentration-independent and serves as a hallmark of gradient sensing. PTEN is a lipid phosphatase that converts PIP_3_ to PIP_2_ and regulates PIP_3_ polarization (Funamoto et al., 2002; Iijima and Devreotes, 2002). The spatiotemporal dynamics of PTEN membrane localization are opposite to those of PIP_3_ in a concentration-dependent manner (Meier-Schellersheim et al., 2006). Upon exposure to low cAMP gradients (<100 nM), cells exhibit a single-phase withdrawal of PTEN from the front and lateral regions, with minimal withdrawal at the rear. In contrast, exposure to high cAMP gradients (>1 μM) elicits a biphasic PTEN response, consisting of an initial uniform withdrawal of PTEN from the entire cell periphery, followed by its return from the cytoplasm to the plasma membrane and then a second withdrawal from the front—representing a typical adaptation process followed by polarized PTEN localization, consistent with PIP_3_ dynamics upon exposure to a steady gradient. Ras activation, an upstream activator of PI_3_K that drives PIP_3_ production, represents the earliest GPCR-mediated signaling step in gradient sensing that exhibits adaptation (Kortholt et al., 2013). Importantly, accumulation of active Ras at the front of gradient-sensing cells depends on gradient concentration. In line with this, we observed concentration-dependent plasma membrane (PM) translocation of C2GAP1 in cells exposed to steady gradients (**Figure 3**). Consistent with preferential C2GAP1 accumulation at the front relative to the back, we previously reported temporally stronger inhibition in the front of PIP_3_-polarized cells in gradients (Xu et al., 2007). Notably, this enhanced inhibition was observed only in cells experiencing strong, high-concentration gradients, but not low-concentration gradients (**Figure 4**). In contrast, *c2gapA⁻* cells failed to exhibit this stronger inhibition in PIP_3_-polarized cells at any gradient concentration, demonstrating an essential role for C2GAP1 in establishing enhanced inhibition in the front during gradient sensing. Agree with the above, *c2gapA^−^* cells display excessive actin polymerization, subsequent broaden leading edge during chemotaxis when experiencing high-concentration gradient, demonstrating the necessity of C2GAP1 accumulation in the leading edge to tune down Ras signaling for proper polarization and efficient cell migration during chemotaxis (Xu et al., 2017). Interestingly, membrane targeting of C2GAP1 does not closely corelate with active Ras on the plasma membrane (Xu et al., 2017) (**Figure 6B**), suggesting the involvement of additional binding partners in C2GAP1 PM localization. In this study, we identify Gα2 as such a plasma membrane binding partner of C2GAP1 in an F-actin–independent manner (**Figure 6A**). Importantly, simulations of binding affinity between C2GAP1 and Gα2 in either the GDP- or GTP-bound state suggest stronger interaction with activated Gα2, providing a mechanism by which Gα2 sustains C2GAP1 on the plasma membrane. Moreover, we detected enhanced G protein activation in *c2gapA⁻* cells (**Figure 6D**), indicating an additional role for the C2GAP1–Gα2 interaction in suppressing G protein activation upon stimulation. This suppressive role of the C2GAP1–Gα2 interaction in regulating G protein activation is essential for rapid reorientation in dynamically changing gradients (**Figure 8**), exemplifying an additional layer of G protein regulation during gradient sensing.

## ACKNOWLEDGMENTS

This research was supported by the Intramural Research Program of the National Institutes of Health (NIH). The contributions of the NIH author(s) are considered Works of the United States Government. The findings and conclusions presented in this paper are those of the author(s) and do not necessarily reflect the views of the NIH or the U.S. Department of Health and Human Services.

## AUTHORS CONTRIBUTIONS

Conceptualization: X.X.; Investigation: X.X., H.H., R.K.; Data analysis: X.X., R.K., H.H., R.D.S.; Writing – Original draft, X.X.; Review & Editing, X.X., H.H., R.K., R.D.S., T.J.; Funding: T.J.

