## supplementary information for "Heterotrimeric G Protein–RasGAP Coupling Drives Adaptation During Chemotaxis"

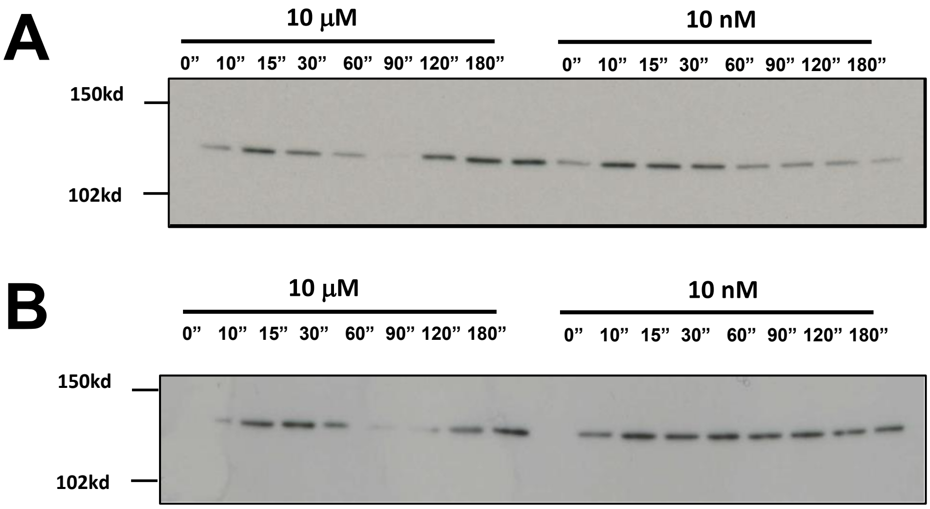
A
10 μM
10 nM
0” 10” 15” 30” 60” 90” 120” 180”
0” 10” 15” 30” 60” 90” 120” 180”
150kd
102kd
B
10 μM
10 nM
0” 10” 15” 30” 60” 90” 120” 180”
0” 10” 15” 30” 60” 90” 120” 180”
150kd
102kd

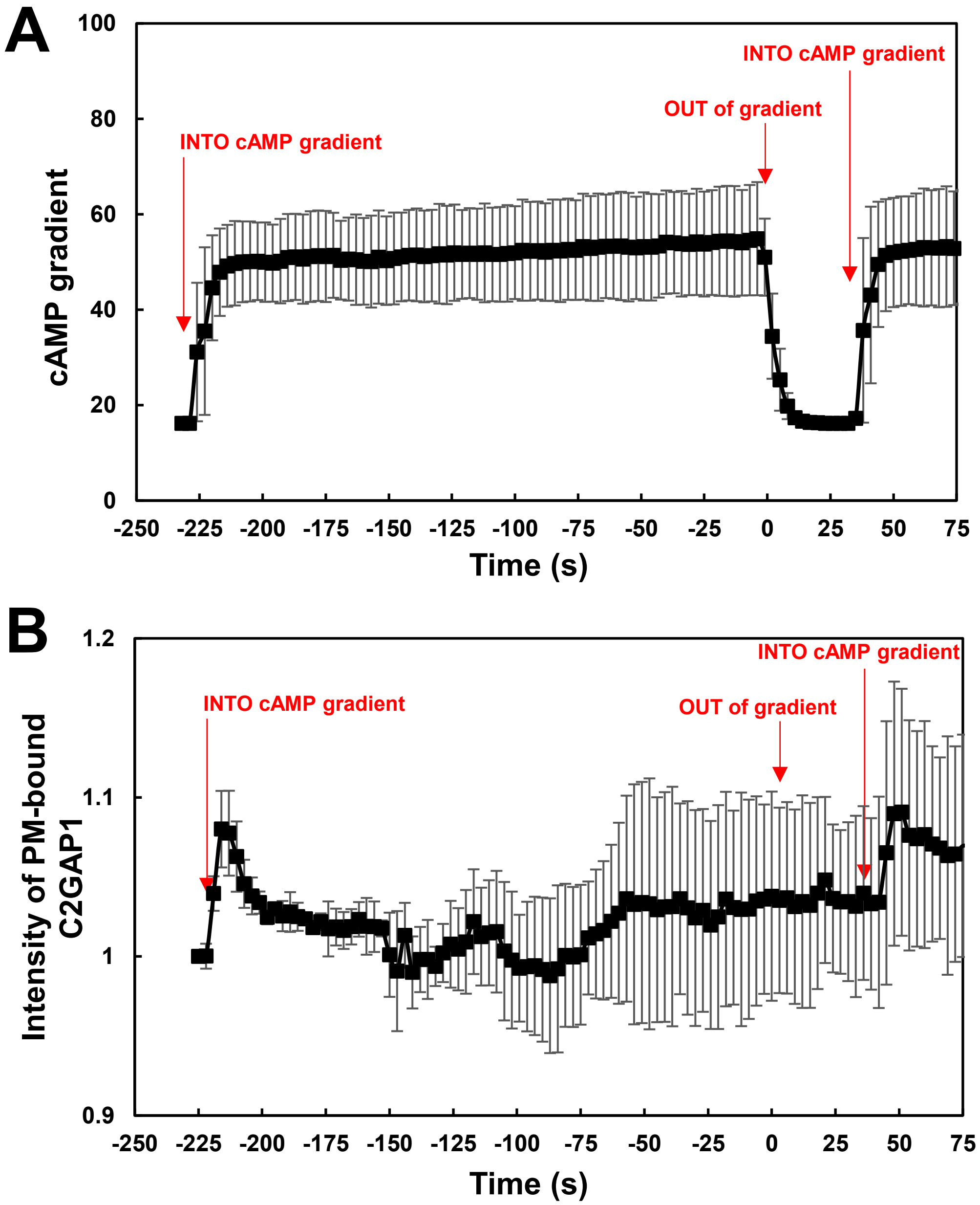
A
100
80
60
40
20
0
cAMP gradient
INTO cAMP gradient
OUT of gradient
INTO cAMP gradient
-250
-225
-200
-175
-150
-125
-100
-75
-50
-25
0
25
50
75
Time (s)
B
1.2
1.1
1
0.9
Intensity of PM-bound C2GAP1
INTO cAMP gradient
OUT of gradient
INTO cAMP gradient
-250
-225
-200
-175
-150
-125
-100
-75
-50
-25
0
25
50
75
Time (s)

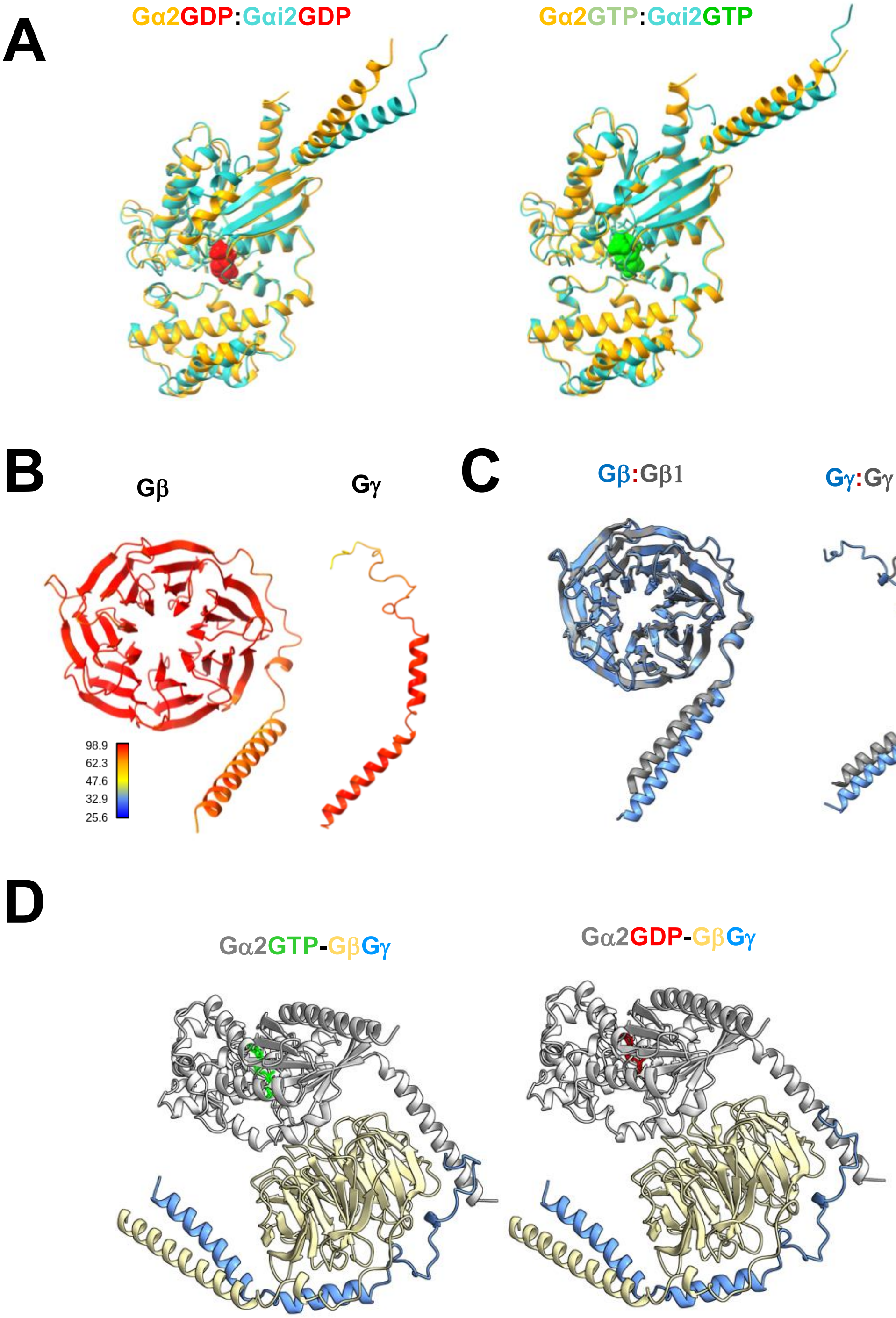
A
Gα2GDP:Gαi2GDP
Gα2GTP:Gαi2GTP
B
Gβ
Gγ
98.9
62.3
47.6
32.9
25.6
C
Gβ:Gβ1
Gγ:Gγ1
D
Gα2GTP-GβGγ
Gα2GDP-GβGγ

**A**

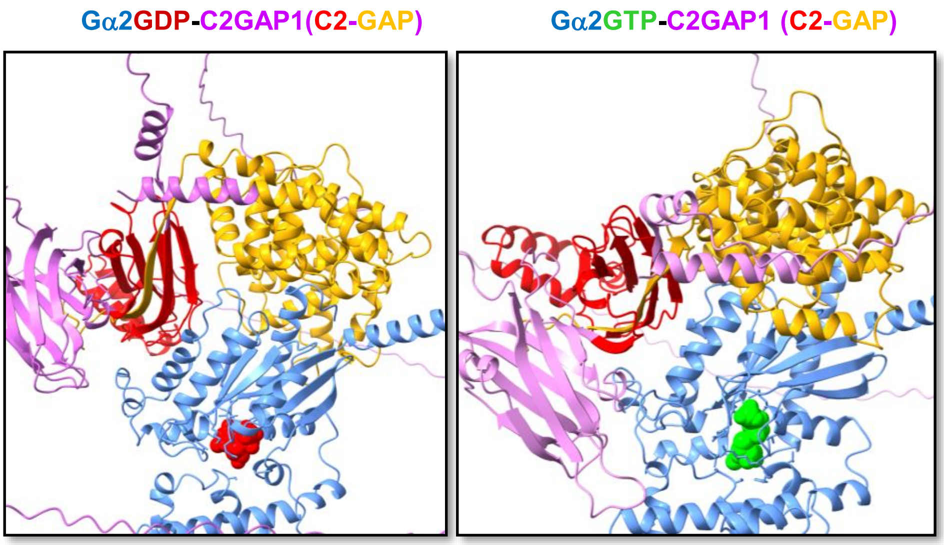


**B**

| | C2GAP1/Gα2-GDP | C2GAP1/Gα2- GTP |
|---|---|---|
| # of intermolecular contacts: | 62 | 73 |
| # of charged contacts: | 14 | 14 |
| # of charged-polar contacts: | 11 | 10 |
| # of charged-apolar contacts: | 17 | 27 |
| # of polar-polar contacts: | 3 | 1 |
| # of apolar-polar contacts: | 6 | 6 |
| # of apolar-apolar contacts: | 11 | 15 |
| % of apolar NIS residues: | 32.21 | 31.90 |
| % of charged NIS residues: | 28.63 | 28.90 |
| Predicted binding affinity (kcal/mol): | -9.8 | -11.2 |
| Predicated dissociation constant: | 6.8e-08 | 6.3e-09 |

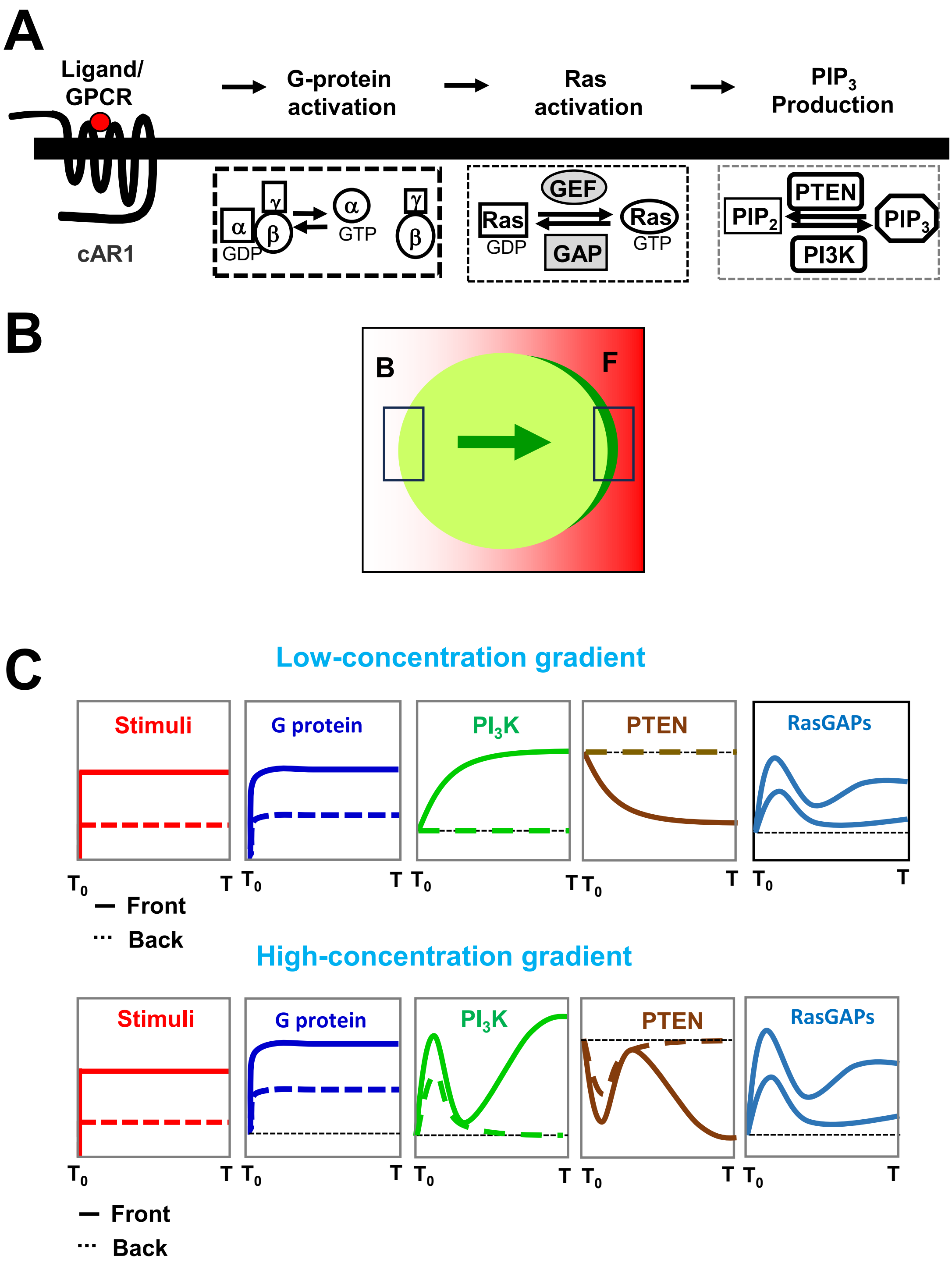
A
Ligand/
GPCR
cAR1
G-protein
activation
α
γ
β
GDP
α
GTP
γ
β
Ras
activation
GEF
Ras
GDP
Ras
GTP
GAP
PIP$_3$
Production
PTEN
PIP$_2$
PIP$_3$
PI3K
B
B
F
C
Low-concentration gradient
Stimuli
G protein
PI$_3$K
PTEN
RasGAPs
T$_0$
T
— Front
··· Back
High-concentration gradient
Stimuli
G protein
PI$_3$K
PTEN
RasGAPs
T$_0$
T
— Front
··· Back

**Supplementary information:**

**Supplemental figure legends:**

**Figure S1.** **Dose-dependent PM targeting C2GAP1 in the cells.** Two independent measurements are shown in **A** and **B** for the quantification of C2GAP1 PM targeting shown in **Figure 3B**.

**Figure S2.** **Dynamics of C2GAP1 in response to application, removal, and reapplication of identical steady gradients.**

**A.** cAMP concentrations during gradient application, removal, and reapplication. cAMP levels were quantified by measuring the fluorescence intensity of Alexa594 surrounding cells; Alexa594 was mixed with cAMP to visualize the gradient.

**B.** Plasma membrane (PM)–associated C2GAP1-YFP in *c2gapA*$^{-}$ cells during gradient application, removal, and reapplication. PM-bound C2GAP1-YFP was quantified as depletion of C2GAP1-YFP from the cytoplasm and normalized to 1 at 0 s. Data are presented as mean ± SD. $N = 5$ and 9 for gradients of 10 μM and 100 nM cAMP, respectively.

**Figure S3** **Predicted structures of heterotrimeric G protein Gα2Gβγ.**

**A.** Structural alignment of *Dictyostelium* Gα2 (gold) and human Gαi2 (cyan) in either GDP-bound (red) or GTP (green)-bound state.

**B.** Predicted structures of *Dictyostelium* Gβ and Gγ with prediction confidence shown by the confidence scale.

**C.** Structural alignment of *Dictyostelium* Gβ (blue) and mammalian Gβ1 (grey), and *Dictyostelium* Gγ (blue) and mammalian Gγ (grey).

**D.** Predicted heterotrimeric G protein complex of *D. discoideum* Gα2Gβγ with Gα2 in either the GDP- or GTP-bound state. Gα2 is shown in grey, Gβ in yellow, and Gγ in blue. GDP and GTP are shown in red and Green, respectively.

**Figure S4.** **Interfaces of Gα2–C2GAP1 interaction.**

**A.** Predicted Gα2–C2GAP1 complexes, showing Gα2 (blue) in GDP-bound (red) or GTP-bound (green) states and the C2 (red) and GAP (gold) domains of C2GAP1 (magenta).

**B.** Predicted parameters of binding affinity between (GDP- or GTP-bound) Gα2 and C2GAP1 interaction.

**Figure S5.** **Distinct dynamics of GPCR cAR1–mediated signaling during gradient sensing in cytoskeleton-free, immobile cells.**

**A.** Four cAR1-mediated signaling events activated by a cAMP gradient during gradient sensing.

**B.** chematic illustrating the front and back regions of a cell, where the signaling dynamics in response to a steady cAMP gradient are quantified in **C**.

**C.** Distinct temporal dynamics of G protein activation and $PIP_3$ production (reflecting $PI_3K$ activation) at the cell front (solid lines) and back (dashed lines) following exposure to either low- or high-concentration cAMP gradients.

**Supplemental videos**

**Video S1.** $PIP_3$ dynamics of gradient sensing in WT (top) or *c2gapA*⁻ (bottom) upon exposure to steady cAMP gradient generated from the source of 100 nM (left) or 10 μM (right) cAMP, respectively. Cells expressing $PIP_3$ biosensor $PH_{Crac}$-GFP (green) were treated with 5 μM latrunculin B 10 min prior to the experiment. To visualize cAMP gradient, cAMP was mixed with fluorescent dye Alexa 594 (red).

**Video S2.** PTEN dynamics of gradient sensing in WT (top) or *c2gapA*⁻ (bottom) upon exposure to steady cAMP gradient generated from the source of 100 nM (left) or 10 μM (right) cAMP, respectively. Cells expressing PTEN-GFP (green) were treated with 5 μM latrunculin B for 10 min prior to the experiment. To visualize cAMP gradient, cAMP was mixed with fluorescent dye Alexa 594 (red).

**Video S3.** C2GAP1-GFP dynamics of gradient sensing in *c2gapA*⁻ upon exposure to steady cAMP gradients generated from the source of either 10 μM (left) or 100 nM (right) cAMP, respectively. Cells expressing C2GAP1-YFP (green) were treated with 5 μM latrunculin B for 10 min prior to the experiment. To visualize cAMP gradient, cAMP was mixed with fluorescent dye Alexa 594 (red).

**Video S4.** $PIP_3$ dynamics of gradient sensing in WT (top) and *c2gapA*⁻ (bottom) cells upon exposure to withdrawal and reapplication of the same cAMP gradient generated from the source of either 10 μM (left) or 100 nM (right) cAMP, respectively. Cells expressing $PIP_3$ biosensor, $PH_{Crac}$-GFP (green) were treated with 5 μM lat. B 10 min prior to the experiment. To visualize cAMP gradient, cAMP was mixed with fluorescent dye Alexa 594 (red).

**Video S5.** C2GAP1-GFP dynamics of gradient sensing in *c2gapA*⁻ cells upon exposure to withdrawal and reapplication of the same cAMP gradient generated from the source of 10 μM cAMP. Cells expressing C2GAP1-YFP (green) were treated with 5 μM latrunculin B for 10 min prior to the experiment. To visualize cAMP gradient, cAMP was mixed with fluorescent dye Alexa 594 (red).

**Video S6.** Migration behavior of WT (left) and *c2gapA*$^{-}$ (right) cells in the gradients generated from the source of either 10 μM (top), 100 nM (middle), or 1 nM (bottom) cAMP, respectively.
